# Diagnosis of Osteoarthritis Subtypes with Blood Biomarkers

**DOI:** 10.1101/366047

**Authors:** Kun Zhao, Junxin Lin, Bingbing Wu, Guofei Sun, Chengrui An, Maswikiti Ewetse Paul, Hongwei Ouyang

## Abstract

**Objective:** To identity osteoarthritis(OA) subtypes with gene expression of peripheral blood mononuclear cells.

**Methods:** Gene expression data (GSE48556) of Genetics osteoARthritis and Progression (GARP) study was downloaded from Gene Expression Omnibus. Principal component analysis and unsupervised clustering were analyzed to identify subtypes of OA and compare major KEGG pathways and cell type enrichment using GSEA and xCell. Classification of subtypes were explored by the utilization of support vector machine.

**Results:** Unsupervised clustering identified two distinct OA subtypes: Group A comprised of 60 patients (56.6%) and Group B had 46 patients (43.3%). A classifier including nine genes and CD4^+^ T cell and Regulatory T cell flow cytometry could accurately distinguish patients from each group (area under the curve of 0.99 with gene expression). Group A is typical degenerative OA with glycosaminoglycan biosynthesis and apoptosis. Group B is related to Graft versus host disease and antigen processing and presentation, which indicated OA has a new type of “Antigen processing and presentation” similarly as that of RA.

**Conclusion:** OA can be clearly classified into two distinguished subtypes with blood transcriptome, which have important significance on the development of precise OA therapeutics.

## INTRODUCTION

Osteoarthritis (OA) is a common degenerative joint disorder characterized by pain, stiffness, limitation of movement and disability^1^, of which pain and disability remain the most common outstanding signs and symptoms^2^. It is results from a combination of risk factors, such as increasing age, gender, obesity, knee malalignment, loading of joints, genetics and low-grade systemic inflammation^1,3,4^. OA affects approximately 18% of women and 10% of men over 60 years of age^5,6^. At the same time, the burden resulting from OA is increasing rapidly with an increase in two major risk factors: ageing population and the epidemics of obesity^7^.

There are currently no disease-modifying osteoarthritis drugs that can be used to treat OA patients. This may be due to the heterogeneity of the OA population, where the disease progression is often poorly understood^8^. Recent researches show that OA has subtypes with joint cartilage transcriptome. Stratification on histopathological features have revealed that the subtypes of OA and new omics technologies have been offered opportunities to analyze different pathogenic mechanisms in the identification of OA subtypes in an unbiased way^9,10^. In order to gain more understanding of OA heterogeneity, several studies have focused on identification of subtypes by a method of DNA methylation and RNA seq data^10,11^. However, they both used cartilage samples because gene expression pattern in cartilage reflects the influence of extrinsic systemic and micro environment, and intrinsic chondrocyte responses^10^, of which is difficult to get and can cause secondary damage. 2013). Increased interest in the development of new diagnostic and prognostic tests for early forms of OA may incorporate the use of blood-based biomarkers^8^.

The use of blood-based liquid biopsies has been recommended to overcome limitations of tissue acquisition in the field of cancer research^12,13^, and the application of blood biomarkers have been used in depression^14^, chronic obstructive pulmonary disease^15^ and Alzheimer disease^16^. Distinctive gene expression signature between OA and normal clients has been detected in peripheral blood mononuclear cells (PBMC) sample^17^. These evidence also can provide the clues for analyzing the subtypes of OA with blood samples.

In this study, we want to analyze the gene expression pattern and identify subtypes of OA which was performed by microarray analysis of the Genetics osteoARthritis and Progression (GARP) study and provide a new diagnostic tool for OA subtypes^17^.

## METHODS

### Data collection

The series matrix file(s) of gene expression data (GSE48556) was download from Gene Expression Omnibus (http://www.ncbi.nlm.nih.gov/geo) which is a case-control study comparing gene expression profiles generated with total messenger RNA obtained from peripheral blood mononuclear cells of controls with that of OA patients from the GARP study^17,18^.The method of study populations selection and RNA isolation were described according to the study of Ramos et al ^17^.

### Analysis strategy

Figure 1 showed progress of the study strategy. Series matrix file(s) data was processed and normalized as previously described^17^. Firstly, using R package to analyze the difference between OA and healthy controls and identity the subtypes of OA population. Then analysis the difference between subtypes and find marker genes to distinguish and to diagnose OA subtypes. Finally, classification and accuracy analysis was taken by the use of machine learning.

**Figure 1.**
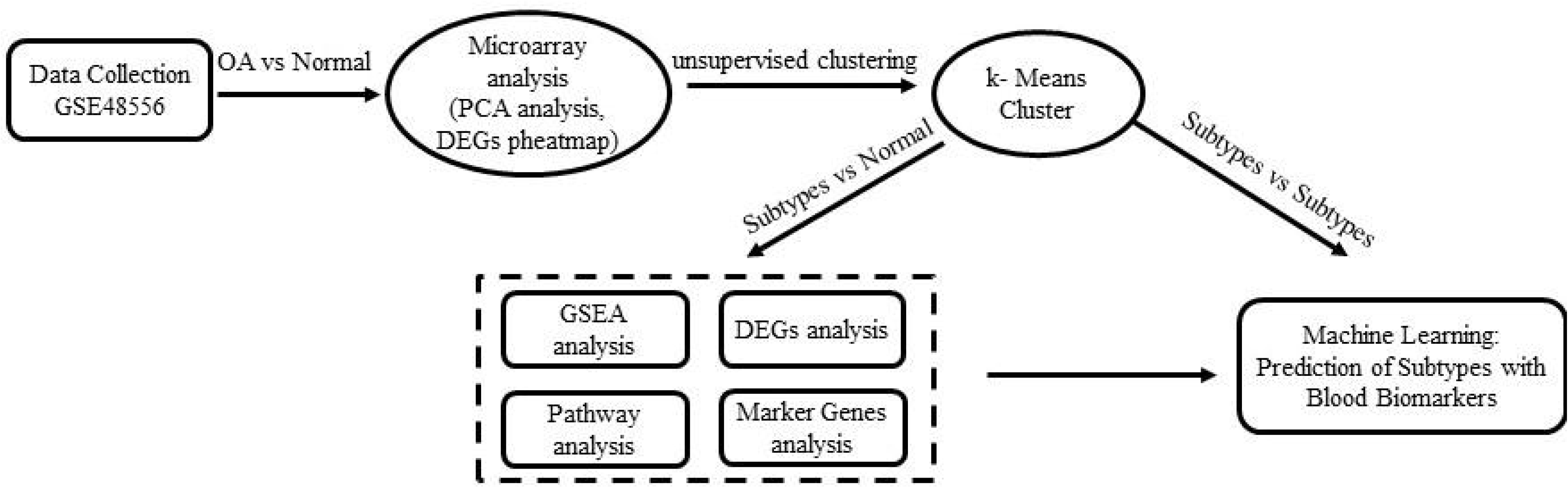

### Microarray analysis

ID conversion was performed by annotate package version 1.58.0 and prcomp function in R 3.4.4 was used to perform principal components analysis(PCA). Differentially expressed genes (DEGs) were identified by limma package version 3.36.1^19^ for linear regression analysis which were reported with Benjamini–Hochberg (BH) corrected p-values of 0.05^19,20^.

### Unsupervised clustering

Unsupervised clustering was carried out by k-means clustering. Using SC3 package version 1.8.0 which was developed as a tool for the unsupervised clustering of single cell RNA-Seq experiments with full microarray gene expression to analysis^21^.The theory of k-means algorithm could also be adapted into the microarray analysis.

### Subtypes identification analysis

Fold changes were calculated with limma package with P values adjusted for multiple testing with Benjamini-Hochberg correction^19^. DEGs were identified with an absolute fold change of 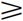 and adjust P value < 0.05. Each subtype identified by the k-means clustering result was compared with healthy controls population to see the difference between subtypes. A support vector machine(SVM) classifier with five-fold cross-validation was performed to distinguish OA subtypes. Models were evaluated by a receiver operating characteristic (ROC) analysis using an area under the curve (AUC) criterion.

### KEGG pathway analysis

Gene set enrichment analysis (GSEA) was performed which used desktop application from the Broad Institute (http://www.broad.mit.edu/gsea/) to do the Kyoto Encyclopedia of Genes and Genomes(KEGG) pathway analysis and FDR q value < 0.05 which were considered to be of significance^22^.

### Analysis of protein interaction networks

To investigate protein interactions among the genes identified in significant KEGG pathway, the search tool for the retrieval of interacting genes/proteins (STRING) 10.5 available online(https://string-db.org/) was used^23^.

### Cell type enrichment analysis

xCell is an online tool for cell type enrichment analysis of 64 immune and stromal cell type with gene expression data (http://xcell.ucsf.edu/)^24^. The tissue cellular heterogeneity landscape of subtypes of OA was used this method to digitally portray.

## RESULTS

### Gene expression pattern in patients with OA and healthy controls

After gene id conversion, 19238 unique genes were presented and PCA analysis visualized the first two principal components and showed huge heterogeneity between OA population (Figure 2A). Analyzing the gene expression pattern by comparing the results of 106 patients with OA to 33 healthy controls, and 1502 DEGs were identified of which adjust P value < 0.05. Different DEGs expression patterns between OA population have been found from heatmap (Figure 2B).

**Figure 2.**
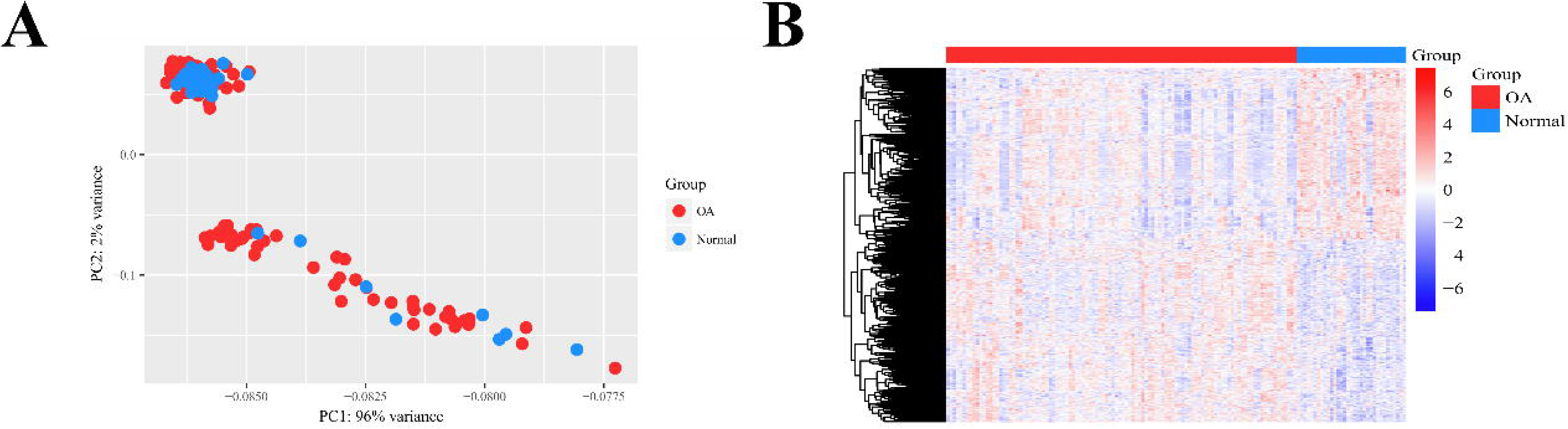

### Stratification of gene expression between two OA clusters

To cluster and identify the subtypes, unsupervised clustering k-means method by SC3 package was used^21^. The most stable clustering distinguished two subgroups and measure of the degree of fit by consensus matrix and silhouette score revealed a strong identity within each cluster (Figure 3A). The gene expression with microarray thus divided the OA patients into two clusters: Group A 56.6% (N=60) and Group B 43.3% (N = 46). To eliminate subtype differences was due to the crowd itself, we also clustered the healthy controls sample, but results showed healthy controls population are homogeneous in which k estimated to be one (Figure 3B). Sixteen differentially expressed genes (DEGs) comparing to the healthy controls population were detected with fold change 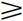 and the adjust P value < 0.05. However, when the results were divided into Group A and Group B and each group was compared separately with the healthy controls population, an additional 550 DEGs were detected, with 11 of those endemic to Group A and 517 distinctive to Group B (Figure 4A). We next want to test whether we can distinguish two main groups by those 11 genes specifically expressed in Group A which ALAS2, CAMP, FAM43A, GPR18, LILRA3 were upregulated and DUSP1, FOS, GNL3L, MAP3K7CL, PDEA4, STK17B were downregulated (Figure 4B), but DUSP1 (P = 0.42) and PDE4C (P = 0.132) did not have significant statistical difference using Student’s t test. The information of nine genes were collected by Gene Cards (http://www.genecards.org/) (Table 1). Then we used to train a SVM classifier with fivefold cross-validation using these nine genes to predict the subtypes with the result of an area under the curve of 0.99 (Figure 4C).

**Figure 3.**
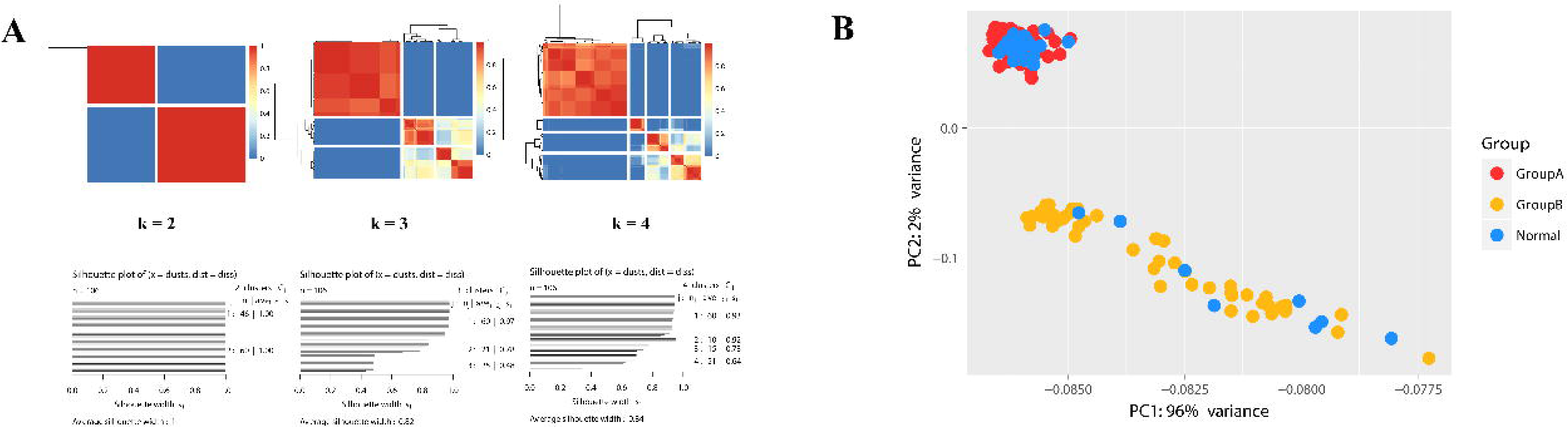

**Figure 4.**
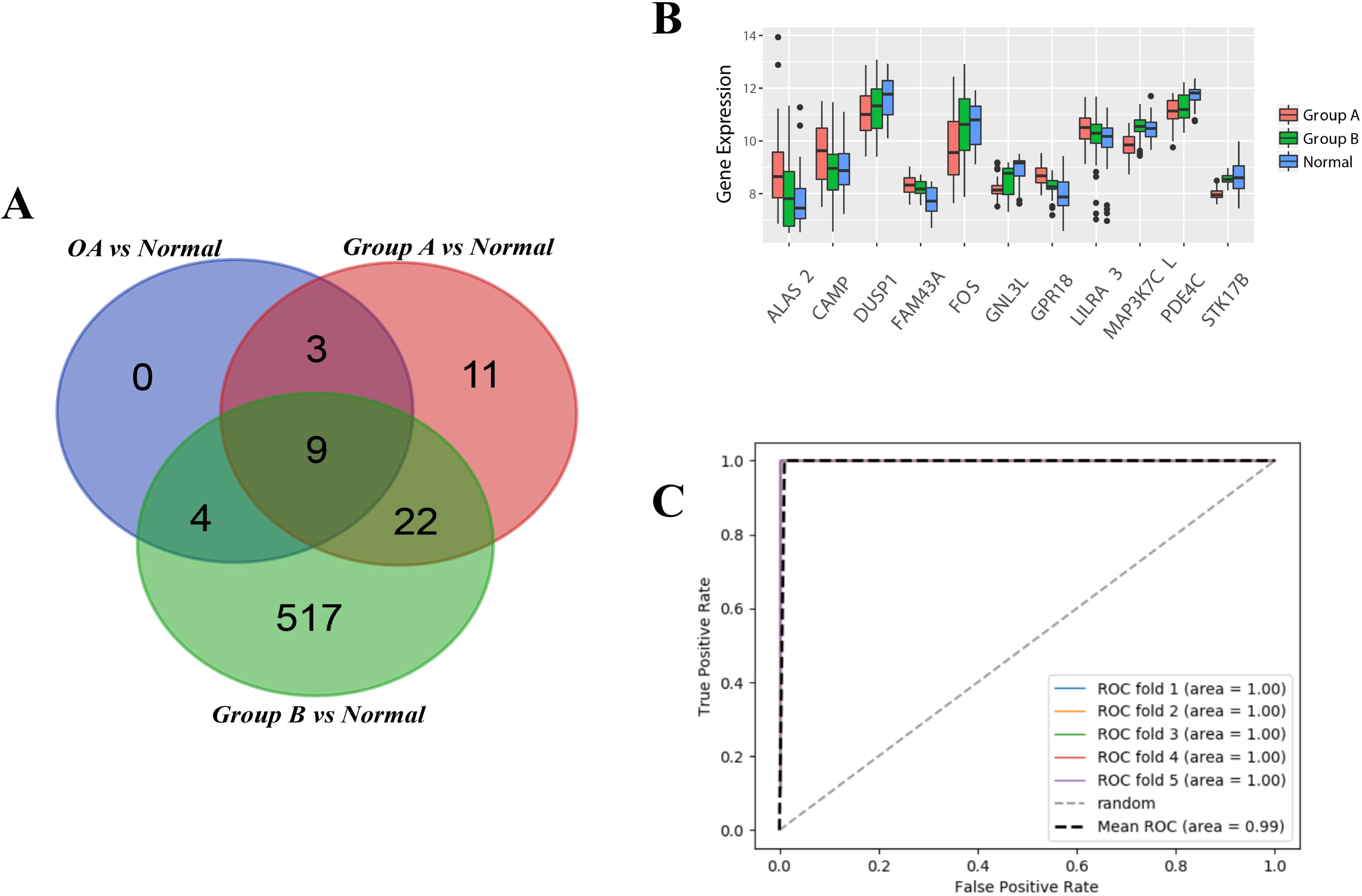

**Table 1.**
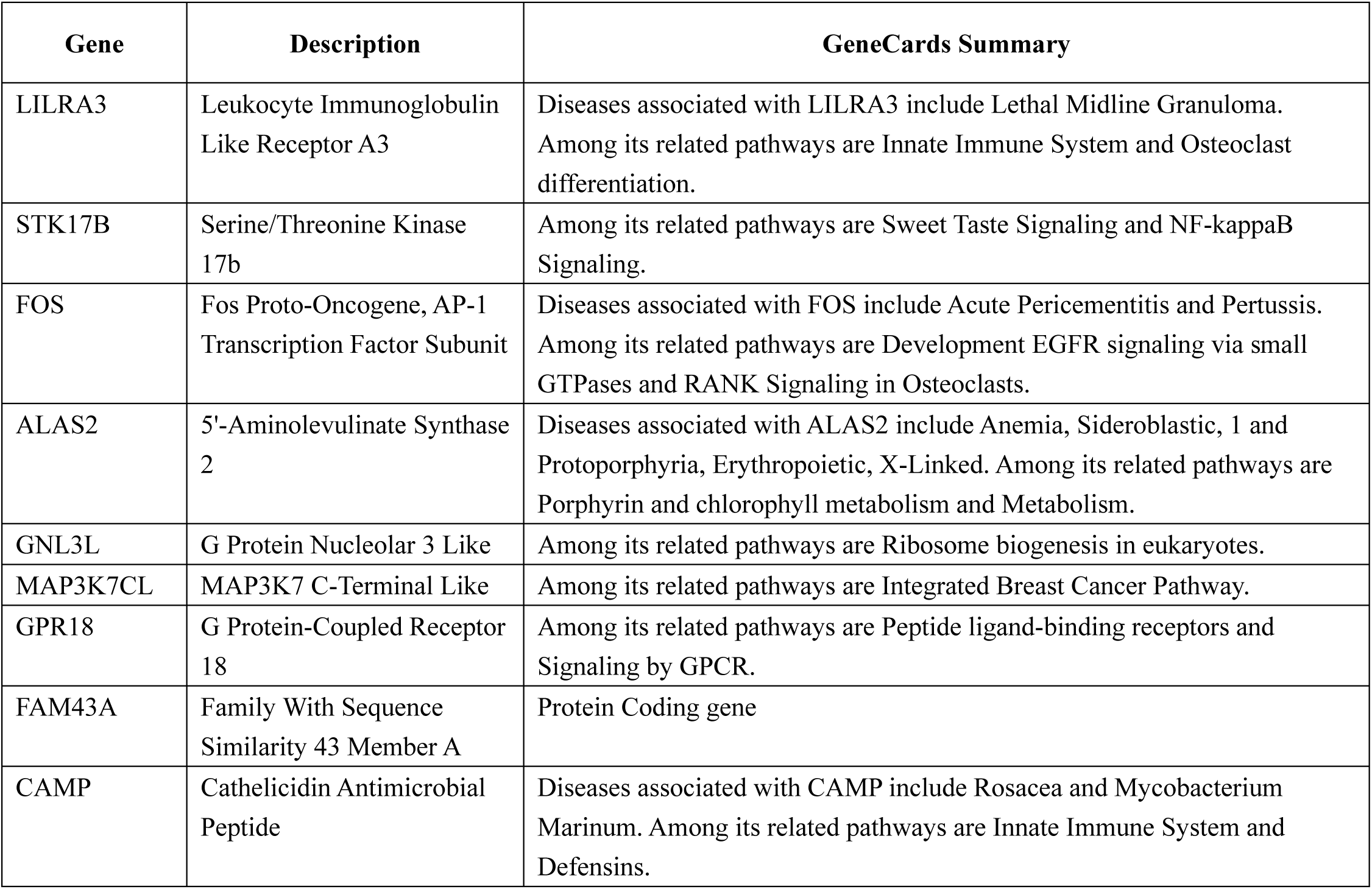
General information of the marker gene distinguish Group A and Group B

### KEGG pathway analysis in subtypes

We then performed GSEA analysis to identify KEGG pathways that are enriched in each group^22^. In contrast to ontology analysis, this algorithm is based on the usage of all gene expression data and derives its function from the analysis of a coordinated sets of genes regulated in a defined biological process or pathway^25^. This identified in Group A changed in Glycosaminoglycan biosynthesis, Porphyrin and chlorophyll metabolism, Glycine serine and threonine metabolism and apoptosis, and Group B changed in Graft versus host disease and Antigen processing and presentation (Figure 5A). Using STRING, we visualized multiple interactions between proteins involved in the significant pathways (Figure 5B).

**Figure 5.**
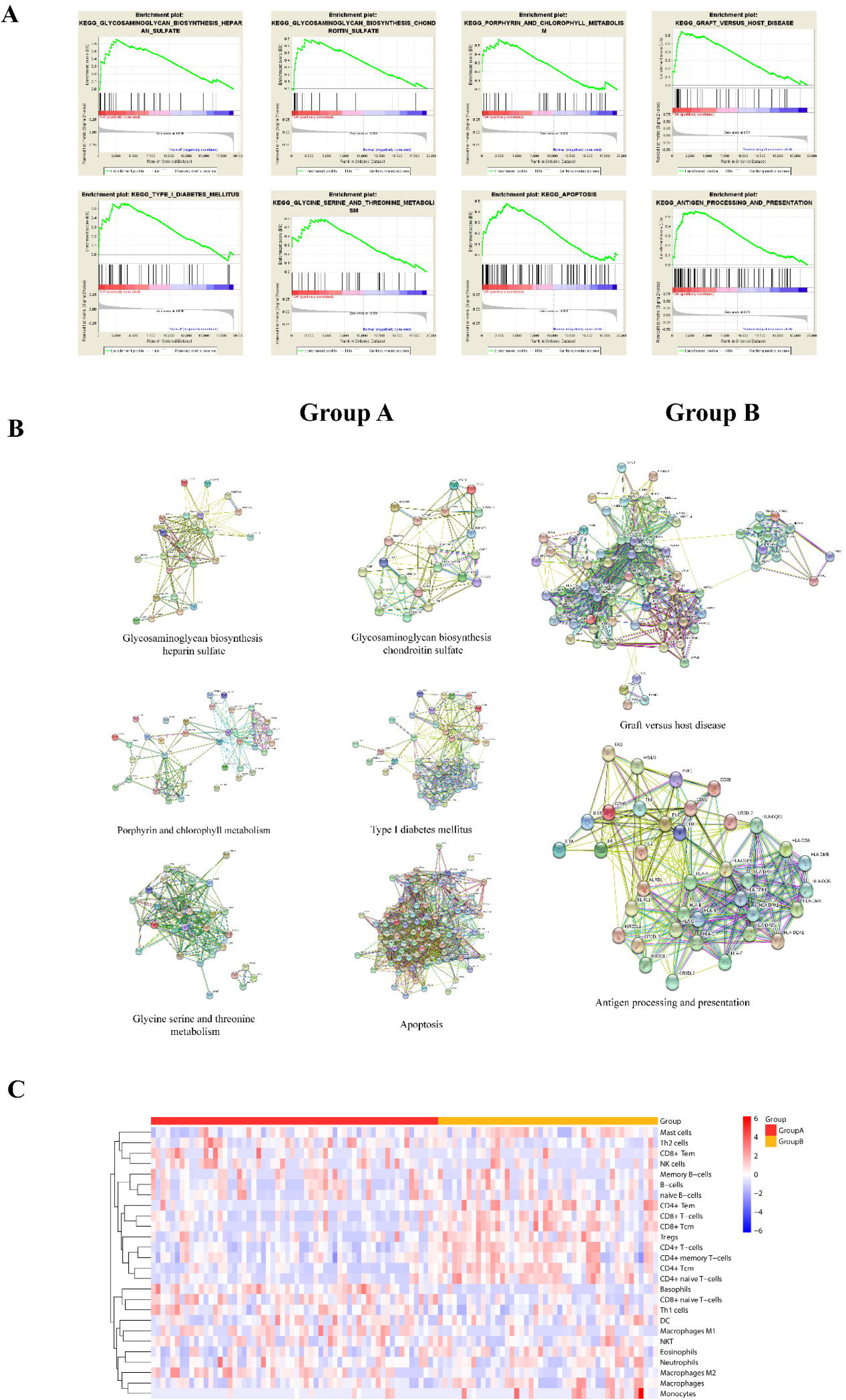

### Cell type enrichment in Group A and Group B

To generate our compendium of gene signatures for cell types, xCell to display cell type enrichments was performed, from which we annotated 106 samples from 64 cell types. We choose 26 out of 64 to see the cell type enrichment pattern. Group B have high relevance with CD4^+^ T Cell, CD4^+^ naive T cells, CD4^+^ Tcm, CD4^+^ Tem, CD8^+^ T Cell, CD8^+^ Tcm and Regulatory T cells (Figure 5C).

## DISCUSSION

OA can be clearly classified into two distinguished subtypes with blood transcriptome, particularly with 9 genes, such as ALAS2, CAMP, FAM43A, GPR18, LILRA3, FOS, CNL3L, MAP3K7CL, STK17B. The two subtypes of OA have distinguished phenotype. One subtype (57%) is typical degenerative OA with glycosaminoglycan biosynthesis and apoptosis. Very interestingly, the other subtype (43%) is related to Graft versus host disease and Antigen processing and presentation through KEGG pathway analysis. Through xCell analysis we further this subtype with the increased number of CD4+ T Cell and Regulatory T cells, which means OA has a new type of “Antigen processing and presentation” similarly as that of RA.

Changes in OA have been implicated in the degenerative process by increasing catabolic activities of metalloproteinase and inhibiting extracellular matrix synthesis^26^. In our study, Group A influenced the glycosaminoglycan biosynthesis which is the important component of extracellular matrix. Our data also hint at the correlation of apoptosis as an aetiological factor for the onset of OA in agreement of the evidence of the study of Ramos et al, but it only affect the Group A^17^.These means Group A is our traditional cognitive osteoarthritis as a degenerative disease.

As for Group B, which is a new type of “Antigen processing and presentation” similarly as that of RA. Some studies have been demonstrated its occurrence not only in rheumatoid arthritis(RA) but also in OA, where it is considered to induce T-cell activation that plays critical roles in autoreactive immune response, and several groups have implicated that T-cells play important roles in the pathogenesis of OA^27-29^. It was interesting that we found relatively overlap in the cell types with the results showing that CD4+ T Cell, CD4+ naive T cells, CD4+ Tcm, CD4+ Tem, CD8+ T Cell, CD8+ Tcm and regulatory T cells were highly enriched. CD4+ T cells are engaged by antigenic peptide fragments in HLA class II molecules cpmplex, and expressed on the surface of professional antigen presenting cells^30^, as the result of HLA-DRA, DRB1, DRB5, DPA1, DPR1, DQA1, DQB1 proteins interactions involved in the “antigen processing and presentation” pathways. This also suggested the activity of DC and macrophages were increased as the number was not change so significant from the result of cell type enrichment. As for the increased number of regulatory T cells which are specialized for immune suppression^31^, maybe due to the number of CD4^+^ T cells in our data. Furthermore, in the field of clinical outcome. Some anti-rheumatic drugs for RA treatment are also effective for OA which means that an antigen-driven immune reaction is a feature of the pathogenesis of OA and in a model of collagen-induced arthritis which type II collagen acts as an autoantigen, cartilage destruction resembles that in OA^29^. It is speculated that chondrocytes have the function of antigen-presenting cells and statistically significant that chondrocytes of OA patients exhibit high expression of DP, DQ, DR, CD80 and CD86 than healthy control group, and mixed culture of T cells and chondrocytes showed that the OA patients with T cell response to chondrocytes were stronger than healthy controls^32^.This suggested when the articular cartilage of OA suffers from chronic degradation damage, the protection provided by the extracellular matrix is also damaged, and the chondrocytes will release type II collagen and activated the autoimmune reaction.

Previous studies have focused on exploring the pathological mechanism of OA, now we have demonstrated that OA have this two main subtypes. We found out that ALAS2, CAMP, FAM43A, GPR18, LILRA3, FOS, CNL3L, MAP3K7CL, STK17B, of these genes expression and the flow cytometry of CD4+ T Cell, CD4+ naive T cells, CD4+ Tcm, CD4+ Tem, CD8+ T Cell, CD8+ Tcm and Regulatory T cells could distinguish subtypes, and it many give some clues for identifying and analyzing the pathological mechanism of OA subtype. As for options for clinical treatment, current treatment can be used such as Non-Steroidal Anti-Inflammatory Drug (NSAID) and hyaluronic acid(HA) for Group A. For Group B, treatment options may be referred to those of managing rheumatoid arthritis that using some disease-modifying anti-rheumatic drugs.

To the best of our knowledge, this is the first paper to utilize blood sample for the analysis of OA subtypes and provide a new insight to research the pathological mechanism of OA. However, this study has some limitations. Our data analysis in based on the GEO data and we cannot get the primary clinical outcomes of the OA patients, so we cannot be sure that the subtype has no association with clinical severity and disease stage. Also, as the limitation of accessing primary data, we cannot adjust the possible batch effects and the result of DEGs between OA and healthy controls which was different form the study of Ramos et al^17^, and the quantitative RT–PCR to validate and replicate the results of the subtype could not either be made use of.

Our study provides a new insight of OA identification and the basic research of OA, but the classifier panel of genes and potential biomarkers in blood requires further validation and testing. Follow-up investigations are needed to confirm potential predictive value, and the result and conclusion need to be replicated and validated by RNA-seq data.

## Acknowledgements

Thank Professor Qin Lin (Department of Orthopedics and Traumatology of Chinese University of Hong Kong) and Wang Lie (School of Medicine of Zhejiang University) for revising the paper. This work was supported by the National Key Research and Development Program of China (2017YFA0104902), NSFC grants (81630065, GZ1094), the Key Scientific and Technological Innovation Team of Zhejiang Province (2013TD11)

## Contributors

All authors have made substantial contributions to the completion of this study. Study conception and design: ZK, Wu BB. Analysis and interpretation of data: ZK, Lin JX and An CR. Classification model: Sun GF. Language polish: Maswikiti Ewetse Paul. Preparation of the manuscript: ZK and Lin JX. Critical reviewing of the manuscript: all authors.

## REFERENCE

1. Martel-Pelletier J, Barr AJ, Cicuttini FM, et al. Osteoarthritis. Nature Reviews Disease Primers. 2016;2:16072.

2. Sheridan C. Low-molecular-weight albumin drug touted for severe osteoarthritis. Nature biotechnology. 2018;36(4):293.

3. Sellam J, Berenbaum F. Is osteoarthritis a metabolic disease? Joint bone spine. 2013;80(6):568–573.

4. Felson DT. The epidemiology of knee osteoarthritis: results from the Framingham Osteoarthritis Study. Paper presented at: Seminars in arthritis and rheumatism1990.

5. Glyn-Jones S, Palmer AJR, Agricola R, et al. Osteoarthritis. Lancet. 2015;386(9991):376–387.

6. Woolf AD, Pfleger B. Burden of major musculoskeletal conditions. Bulletin of the World Health Organization. 2003;81(9):646–656.

7. Bitton R. The economic burden of osteoarthritis. The American journal of managed care. 2009;15(8 Suppl):S230–235.

8. Bay-Jensen A-C, Henrotin Y, Karsdal M, Mobasheri A. The need for predictive, prognostic, objective and complementary blood-based biomarkers in osteoarthritis (OA). EBioMedicine. 2016;7:4–6.

9. Waarsing JH, Bierma-Zeinstra SM, Weinans H. Distinct subtypes of knee osteoarthritis: data from the Osteoarthritis Initiative. Rheumatology. 2015;54(9):1650–1658.

10. Soul J, Dunn SL, Anand S, et al. Stratification of knee osteoarthritis: two major patient subgroups identified by genome-wide expression analysis of articular cartilage. Annals of the rheumatic diseases. 2017:annrheumdis-2017-212603.

11. Fernández-Tajes J, Soto-Hermida A, Vázquez-Mosquera ME, et al. Genome-wide DNA methylation analysis of articular chondrocytes reveals a cluster of osteoarthritic patients. Annals of the rheumatic diseases. 2014;73(4):668–677.

12. Crowley E, Di Nicolantonio F, Loupakis F, Bardelli A. Liquid biopsy: monitoring cancer-genetics in the blood. Nature reviews Clinical oncology. 2013;10(8):472.

13. Best MG, Sol N, Kooi I, et al. RNA-Seq of tumor-educated platelets enables blood-based pan-cancer, multiclass, and molecular pathway cancer diagnostics. Cancer cell. 2015;28(5):666–676.

14. Belzeaux R, Formisano-Tréziny C, Loundou A, et al. Clinical variations modulate patterns of gene expression and define blood biomarkers in major depression. Journal of psychiatric research. 2010;44(16):1205–1213.

15. Kim DK, Cho MH, Hersh CP, et al. Genome-wide association analysis of blood biomarkers in chronic obstructive pulmonary disease. American journal of respiratory and critical care medicine. 2012;186(12):1238–1247.

16. Jellinger KA, Janetzky B, Attems J, Kienzl E. Biomarkers for early diagnosis of Alzheimer disease:‘ALZheimer ASsociated gene’-a new blood biomarker? Journal of cellular and molecular medicine. 2008;12(4):1094–1117.

17. Ramos YF, Bos SD, Lakenberg N, et al. Genes expressed in blood link osteoarthritis with apoptotic pathways. Annals of the rheumatic diseases. 2013:annrheumdis-2013-203405.

18. Riyazi N, Meulenbelt I, Kroon HM, et al. Evidence for familial aggregation of hand, hip, and spine but not knee osteoarthritis in siblings with multiple joint involvement: the GARP study. Annals of the rheumatic diseases. 2005;64(3):438–443.

19. Ritchie ME, Phipson B, Wu D, et al. limma powers differential expression analyses for RNA-sequencing and microarray studies. Nucleic acids research. 2015;43(7):e47-e47.

20. Smyth GK. Linear models and empirical bayes methods for assessing differential expression in microarray experiments. Statistical applications in genetics and molecular biology. 2004;3(1):1–25.

21. Kiselev VY, Kirschner K, Schaub MT, et al. SC3: consensus clustering of single-cell RNA-seq data. Nature methods. 2017;14(5):483.

22. Subramanian A, Tamayo P, Mootha VK, et al. Gene set enrichment analysis: a knowledge-based approach for interpreting genome-wide expression profiles. Proceedings of the National Academy of Sciences. 2005;102(43):15545–15550.

23. Szklarczyk D, Morris JH, Cook H, et al. The STRING database in 2017: quality-controlled protein-protein association networks, made broadly accessible. Nucleic acids research. 2016:gkw937.

24. Aran D, Hu Z, Butte AJ. xCell: Digitally portraying the tissue cellular heterogeneity landscape. Genome biology. 2017;18(1):220.

25. Van der Pouw Kraan T, Wijbrandts C, Van Baarsen L, et al. Rheumatoid arthritis subtypes identified by genomic profiling of peripheral blood cells: assignment of a type I interferon signature in a subpopulation of patients. Annals of the rheumatic diseases. 2007;66(8):1008–1014.

26. Rahmati M, Mobasheri A, Mozafari M. Inflammatory mediators in osteoarthritis: A critical review of the state-of-the-art, current prospects, and future challenges. Bone. 2016;85:81–90.

27. Revell PA, Mayston V, Lalor P, Mapp P. The synovial membrane in osteoarthritis: a histological study including the characterisation of the cellular infiltrate present in inflammatory osteoarthritis using monoclonal antibodies. Annals of the Rheumatic Diseases. 1988;47(4):300.

28. Sakkas LI, Scanzello C, Johanson N, et al. T cells and T-cell cytokine transcripts in the synovial membrane in patients with osteoarthritis. Clinical and diagnostic laboratory immunology. 1998;5(4):430–437.

29. Ishii H, Tanaka H, Katoh K, Nakamura H, Nagashima M, Yoshino S. Characterization of infiltrating T cells and Th1/Th2-type cytokines in the synovium of patients with osteoarthritis. Osteoarthritis and cartilage. 2002;10(4):277–281.

30. Ulvestad E, Williams K, Bo L, Trapp B, Antel J, Mork S. HLA class II molecules (HLA-DR, -DP, -DQ) on cells in the human CNS studied in situ and in vitro. Immunology. 1994;82(4):535–541.

31. Sakaguchi S, Yamaguchi T, Nomura T, Ono M. Regulatory T cells and immune tolerance. Cell. 2008;133(5):775–787.

32. Alsalameh S, Jahn B, Krause A, Kalden J, Burmester G. Antigenicity and accessory cell function of human articular chondrocytes. The Journal of rheumatology. 1991;18(3):414–421.

